# Generating an efficient arginase variant for medical and industrial uses: In Silico engineering

**DOI:** 10.1101/2023.03.27.534378

**Authors:** Haitham Ahmed Al-Madhagi, Mahdi H. Alsugoor

**Author notes:** **Correspondence**: Haitham Ahmed Al-Madhagi, Biochemical Technology Program, Dhamar University, Yemen.

## Abstract

Human arginase is a multifaceted enzyme that can be utilised for various medical and industrial applications, including as a replacement therapy for enzyme-deficient patients and for the industrial production of ornithine. However, no report has explored the in-silico engineering of this novel enzyme. The crystal structure of human arginase 1 was downloaded from the protein databank, and its quality was checked prior to further analysis. CUPSAT and DeepDDG webservers were then employed to nominate the most stable variants, which were prepared by the UCSF Chimera v1.16 modelling system and refined by the GalaxyRefine tool. Docking (i.e., to reference substrate and inhibitor), stability confirmation and dynamics simulations were conducted for all proposed variants, compared to the wild-type version of the enzyme. G119L was the best mutant in all the mentioned aspects, which was afterwards cloned in silico as a final step for the experimental testing thereof. Accordingly, G119L is found to be a valuable arginase mutant that deserves experimental validation to be employed for medical and industrial purposes.

## Introduction

L-arginine is a basic, semi-essential amino acid (AA) required for the synthesis of other AA proteins and polyamines. Arginine is considered a semi-essential AA, because it can be synthesised *de novo*, but cannot satisfy the required amount, especially during childhood. It plays an important role in detoxifying ammonia, nitric oxide synthesis, immunomodulation and hormonal secretion [1]. In the past decade, arginine has been used as an important supplement for muscle growth through nitric oxide-mediated vasodilation of endothelium, which increases nutrient support to striated muscles and eliminating muscle metabolites after training [2]. Moreover, arginine is one of the metabolites in the urea cycle purposed for ammonia detoxification; it is broken down via the arginase enzyme to urea, the less-toxic form of ammonia ready for excretion; and ornithine, which is necessary for cellular growth stimulation. Arginase (EC 3.5.3.1) is a heterotrimeric metalloenzyme with Mn^+2^, a cofactor in each subunit [3]. Arginase is a ubiquitous enzyme present in prokaryotes, yeasts, plants, invertebrates and vertebrates and contributes to critical division and development of those organisms [4]. In humans, arginase has two isozymes: arginase 1 is 322 and arginase 2 is 354 AA residues encoded by different genes in distinct chromosomes. They share nearly 50% sequence homology, and this percentage increases to 100% in the catalytic region [5].

Arginase has been shown to have potential for application in distinct aspects, including cardiovascular (CVS) and cancer therapy as a replacement for enzyme-deficient patients and for the industrial production of ornithine [6]. In the general population, it was estimated that CVS deaths accounted for more than 50% of men and approximately 44% of women, compared with nearly 10% and 8%, respectively, in the cancer cohorts [7]. Additionally, the survivors of cancers have a medium-to long-term risk of developing CVS diseases compared to the normal population [8]. Accumulating evidence suggests the implication of arginase in atherosclerosis [9], hypertension [10] and ischemic reperfusion (IR) [11]; arginase was also reported to be upregulated in these conditions and in certain types of cancer [12]. Conversely, arginase-2 protected myocardial IR injury in rats by suppressing the nuclear factor kappa B/tumour necrosis factor (NF-κB/TNF-α) pathway and thereby blocking the inflammatory response [13]. Arginase is administered to treat arginine auxotrophic tumours by depleting the highly demanded arginine and thus brakes the cancer tissue division and proliferation [14].

Arginase deficiency is a rare genetic condition that results in hyperargininemia and subsequent developmental retardation, intellectual disability and mobility defects [15]. Current management is protein restriction, essential AA supplementation and ammonia scavenger’s; despite these precautions, some children die before reaching adolescence due to severe metabolic decompensation [16]. Although a liver transplant alleviates metabolic and biochemical problems, neurologic and mobility defects remain uncontrollable [17]. While no licensed therapeutic for this enzyme deficiency has been approved to date, a clinical trial experimenting PEGzilarginase as a replacement therapy is being undertaken [18]; this forces medical care practitioners to only focus on controlling arginine levels to manage this rare condition [19].

Ornithine is a non-essential, non-protein AA that is found in meats, fish, dairy and eggs. Once ingested, its main role is participation in the urea cycle. In addition, ornithine is known for its potency as an agonist for the AA receptor G-protein-coupled receptor family C group 6 subtype A (GPRC6A) and calcium-sensing receptors providing an extraordinary flavour action [20,21]. L-ornithine—but not D-isomer—was employed to enhance preference to sweet, salty, umami and fat taste but not as a flavour [22]; this substitutes the commonly used counterpart, i.e. monosodium glutamate since many reports confirmed the adverse effects thereof, especially in excessive amounts [23,24]. Such a natural flavour is synthesised from arginine via arginase enzyme through metabolically engineered bacterial fermentation [25].

These facts prioritise the preliminary engineering of arginase enzymes through bioinformatics tools to be used for the medical as well as industrial applications, which is the goal of this study.

## Methods

The planned strategy for the execution of this current work is summarised in Fig. 1.

**Figure 1.**
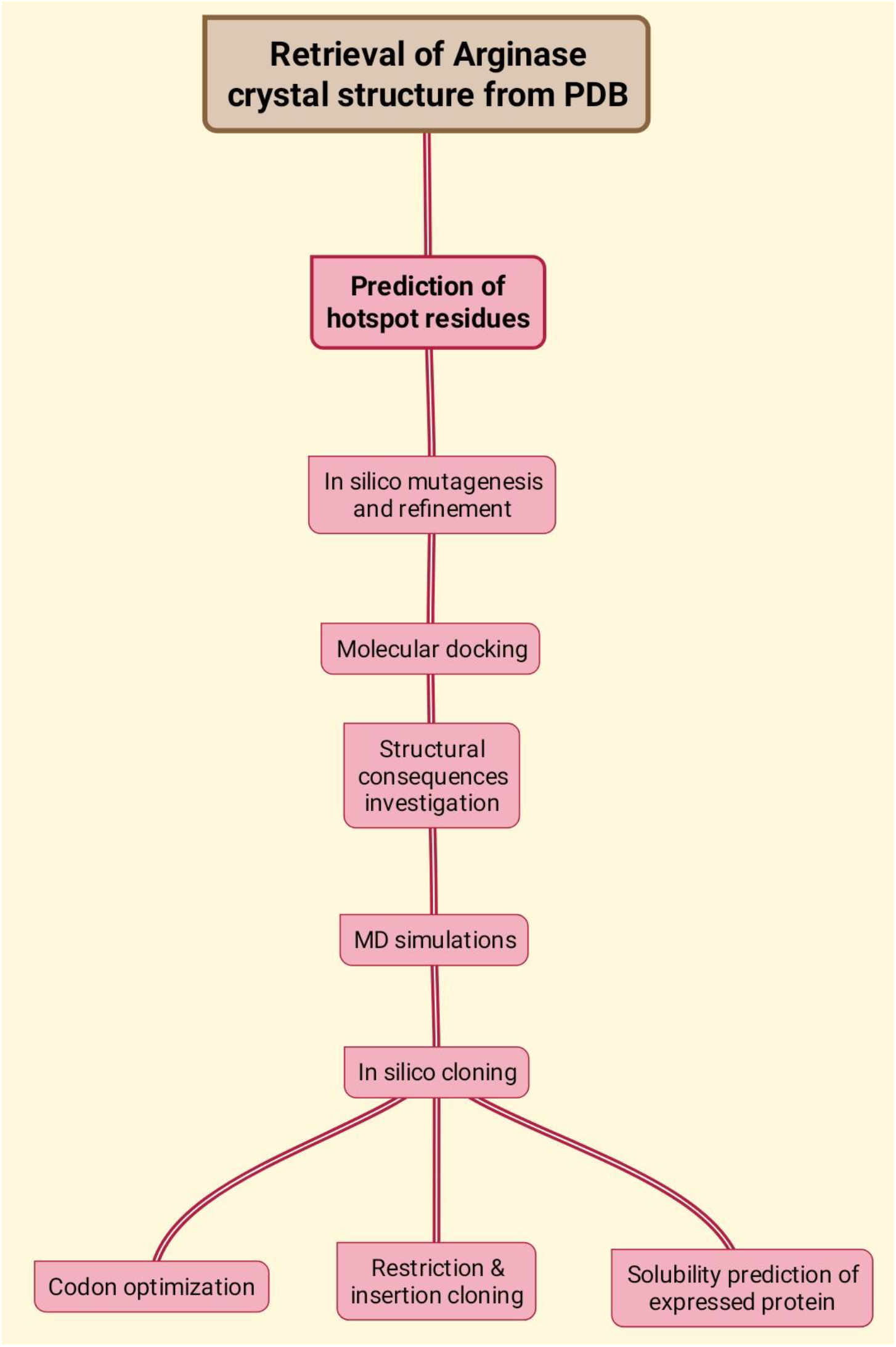
Planned outline of workflow to accomplish the current study.

### Retrieval and check of the tertiary structure of arginase

Human arginase 1 was downloaded from the protein data bank (PDB code: 2AEB). The crystal structure had a resolution of 1.29 Å [26]. All heteroatoms and water molecules were removed. Only the chain A was considered for further exploration. The quality of the prepared protein tertiary structure was assessed using the SWISS-MODEL [27] and ProsA tools [28].

### Prediction of hotspot AA residues

CUPSAT [29] and DeepDDG webservers [30] were utilised to rank the top stabilising and enhancing mutations that would improve enzyme binding to its native substrate. In the two webservers, catalytic residues were excluded from in silico mutagenesis.

### In silico mutagenesis

UCSF Chimera v1.16 was employed for the purpose of in silico mutagenesis. The predicted hotspot residues and the corresponding mutants were prepared, and the highest probable rotamers of the mutant conformers were chosen. This was followed by the energy minimisation and refinement of the produced mutant via the GalaxyRefine tool [31]. Models with the lowest MolProbity, clashes and highest Ramachandran-favoured percentage were selected for the next step.

### Molecular docking

The refined models of the generated mutants were subjected to molecular docking through the CB-Dock-2 tool [32]. They were docked to both reference substrate and inhibitor, keeping all options as default. Pocket 1 was selected for all docking processes. The docked complex was investigated for the interactions formed during docking using the Discovery Studio Client 2021 program.

### Structural effects of mutation

To evaluate the impact of the generated mutation on the 3D architecture of the enzyme, the PremPS server [33] was used. In addition to revealing the detailed interaction network of the mutants, PremPS server also predict the impact of such mutation on the overall stability of the enzyme structure.

### Molecular dynamics simulations

Molecular dynamics (MD) simulations were conducted employing the WebGro webserver [34]. GROMOS96 43a1 was the selected forcefield, the water model was SPC, the box type was triclinic, and the salt type was NaCl with neutralisation and at 0.15 M concentration. The energy was minimised through steepest descent with 5,000 steps. The equilibration type was NVT/NPT at 300 K, 1 bar. MD simulation was conducted for 50 ns to determine the relative-mean square deviation (RMSD), relative mean square fluctuations (RMSF), radius of gyration (Rg) and solvent-accessible surface area (SASA).

### In silico cloning

The mutant that yielded the best binding affinity towards arginine was chosen for the in-silico cloning process. The protein sequence was back-translated by the EMBOSS: backtranseq tool, then the sequence output was uploaded to the JCat tool [35]; this tool performs codon optimisation based on the expressing system in order to maximally increase the likelihood of gene expression in that system. Afterwards, the optimised sequence was inserted to pET-28a(+) vector through SnapGene 6.2 software. The gene was inserted between EcoRI and BamHI restriction enzyme sites. Finally, the solubility of the predicted gene to be translated was assessed via the Protein-sol tool [36].

## Results

### Quality assessment

MolProbity and clash scores of the WT and G119L tertiary structures were 1.21, 1.30 and 1.30, 5.59, respectively, and the Ramachandran-favoured percentage was the same (98.40%). Otherwise, the Ramachandran plot, ProsA Z scores (−8.79 vs. −8.66) and energy profiles were not significantly different, as shown in Fig. 2.

**Figure 2.**
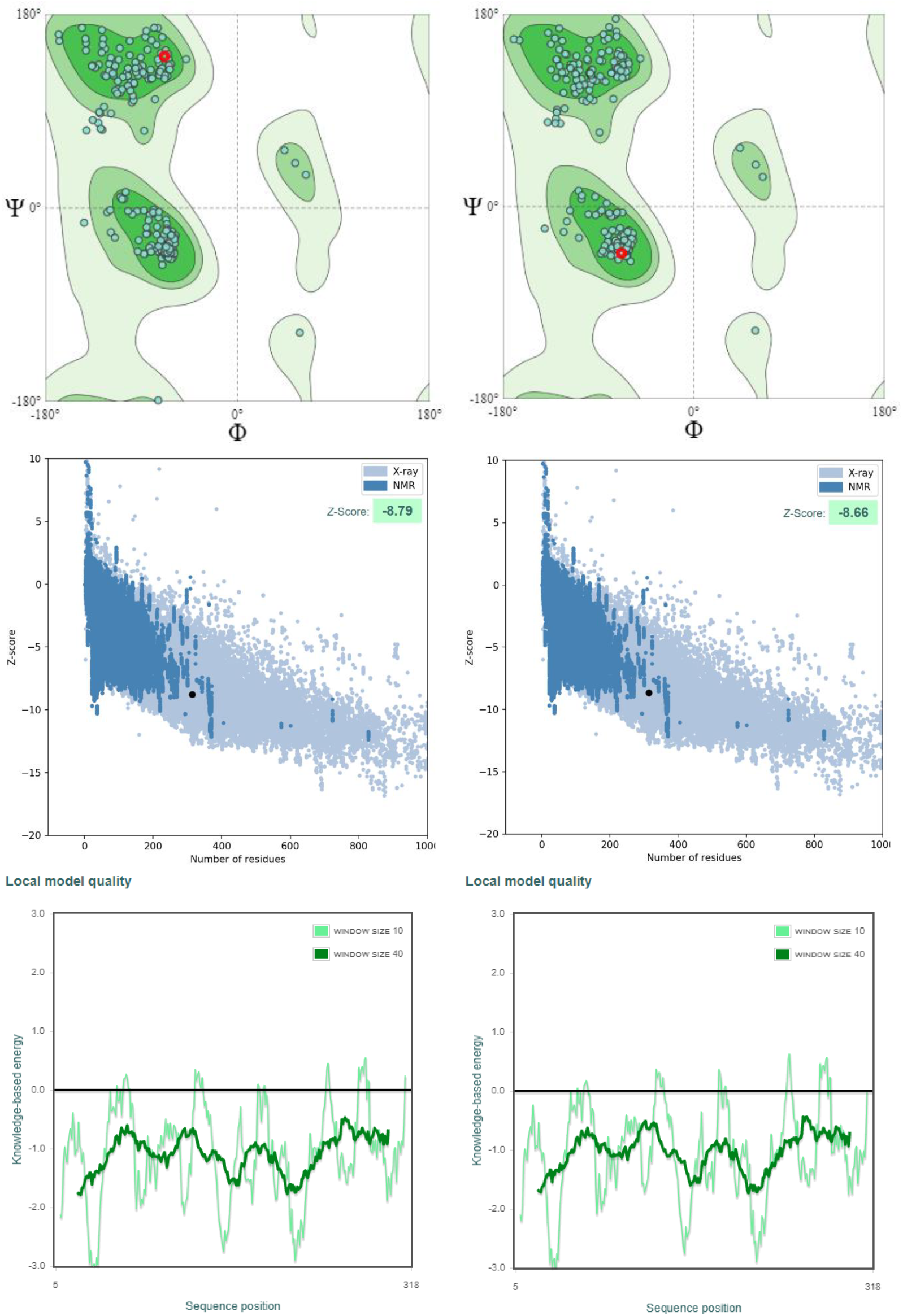
Quality assessment of wild-type arginase and G119L using Ramachandran plot and ProsA and energy plots.

### Molecular docking

Sixteen mutants were tested for their potential to enhance enzyme activity in terms of binding the native substrate relative to the WT version. Among the top-ranked potential variants in CUPSAT and DeepDDG, G119L and, to a lesser extent, D274I have a higher affinity towards the substrate arginine (−5.9 vs. −5.6 kcal/mol). Moreover, G119L had slightly weaker binding affinity towards the reference inhibitor (Table 1). The G119L mutant was therefore considered for further analysis.

**Table 1.** Docking scores to reference substrate and inhibitor.

| Enzyme | Substrate affinity | Inhibitor affinity |
| --- | --- | --- |
| WT | −4.6 | −5.6 |
| G119L | −5.9 | −5.4 |
| G272L | −5.2 | −5.3 |
| K88L | −4.3 | −5.6 |
| S94P | −5.1 | −5.6 |
| D274I | −5.6 | −5.9 |
| D158T | −4.3 | −5.5 |
| G154V | −4.2 | −5.2 |
| I156R | −4.1 | −5.0 |
| K153E | −5.1 | −5.2 |
| P160W | −5.4 | −5.2 |
| S163N | −3.8 | −4.7 |
| V165H | −3.6 | −5.0 |
| G142L | −2.6 | −2.9 |
| P28L | −3.6 | −4.3 |
| P70L | −4.4 | −4.1 |
| S230C | −4.5 | −3.7 |

WT: wild-type

### Mutant impact on enzyme structure

As shown in Fig. 3, the G119L mutant is more powerful than the WT, as reflected by the numerous interactions (i.e., hydrophobic and polar H-bonds) with the surrounding residues near the catalytic pocket. WT only formed two H-bonds with His 228 and Pro 226, while the G119L variant formed two H-bonds with Leu 227 and His 228, in addition to numerous hydrophobic interactions with Ile 121, Leu 219, Leu 220, Arg 225 and Leu 227. This is traced to the bulky side chain of Leu in comparison with Gly, which only has hydrogen in its side chain. Concerning the deleteriousness of the mutation on enzyme stability, the PremPS score was −0.12, which indicates the stabilising effect of G119L on the overall enzyme architecture.

**Figure 3.**
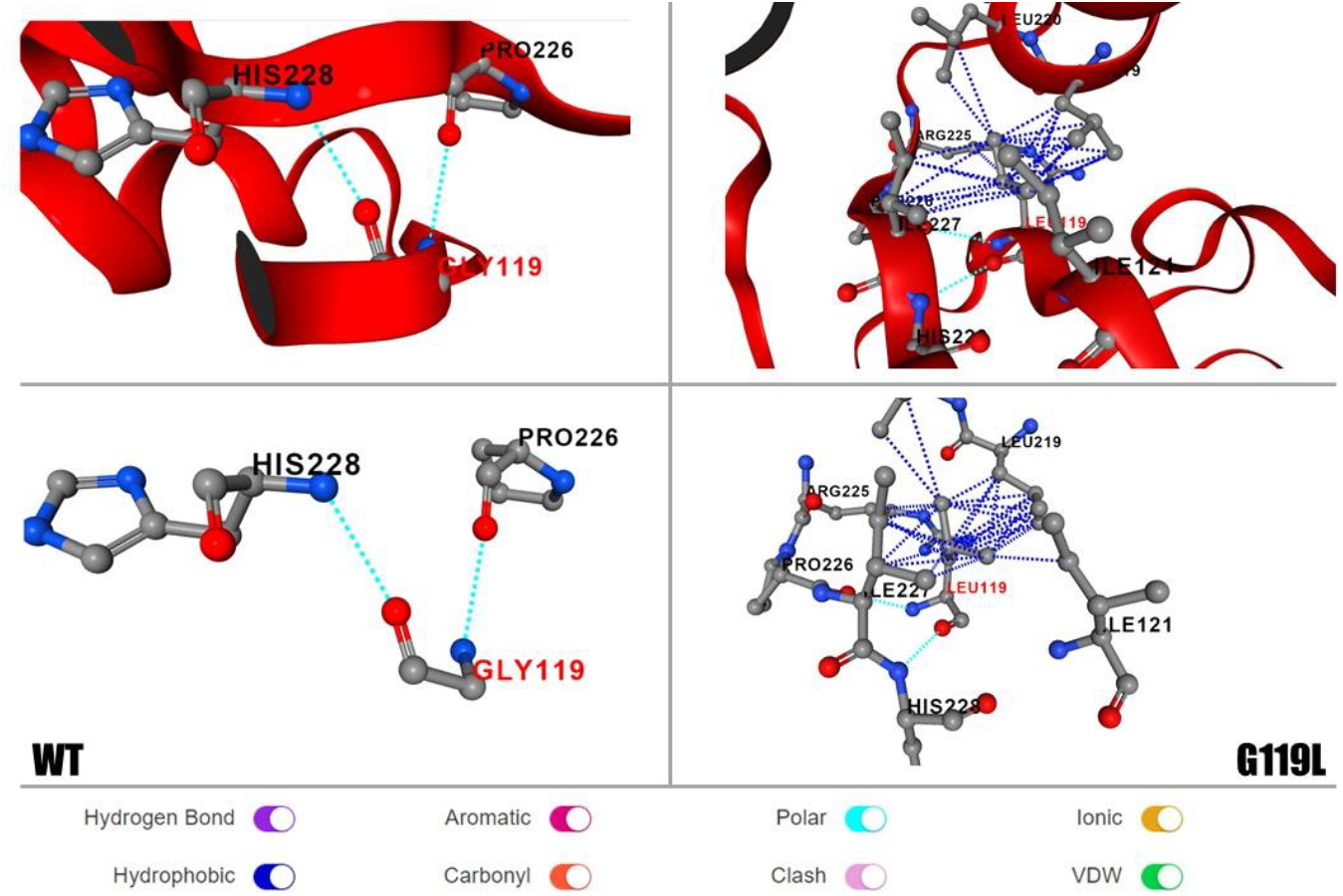
Structural consequence of the G119L mutant (right) compared to WT enzyme (left).

### Protein-ligand interaction

The docked complex (i.e., G119L-arginine) is elucidated in Fig. 4, which illustrates the fit of the substrate in the active centre of the engineered enzyme. However, this binding accounts for the five hydrogen bonds formed between Asp 128, Thr 136 and Ser 137 and the two bonds between Glu 186 and the native substrate. Additionally, salt bridges and Pi-Cation interactions significantly contributed to the strong bonding; these explain the strong binding affinity of the G119L variant to arginine (−5.9 kcal/mol).

**Figure 4.**
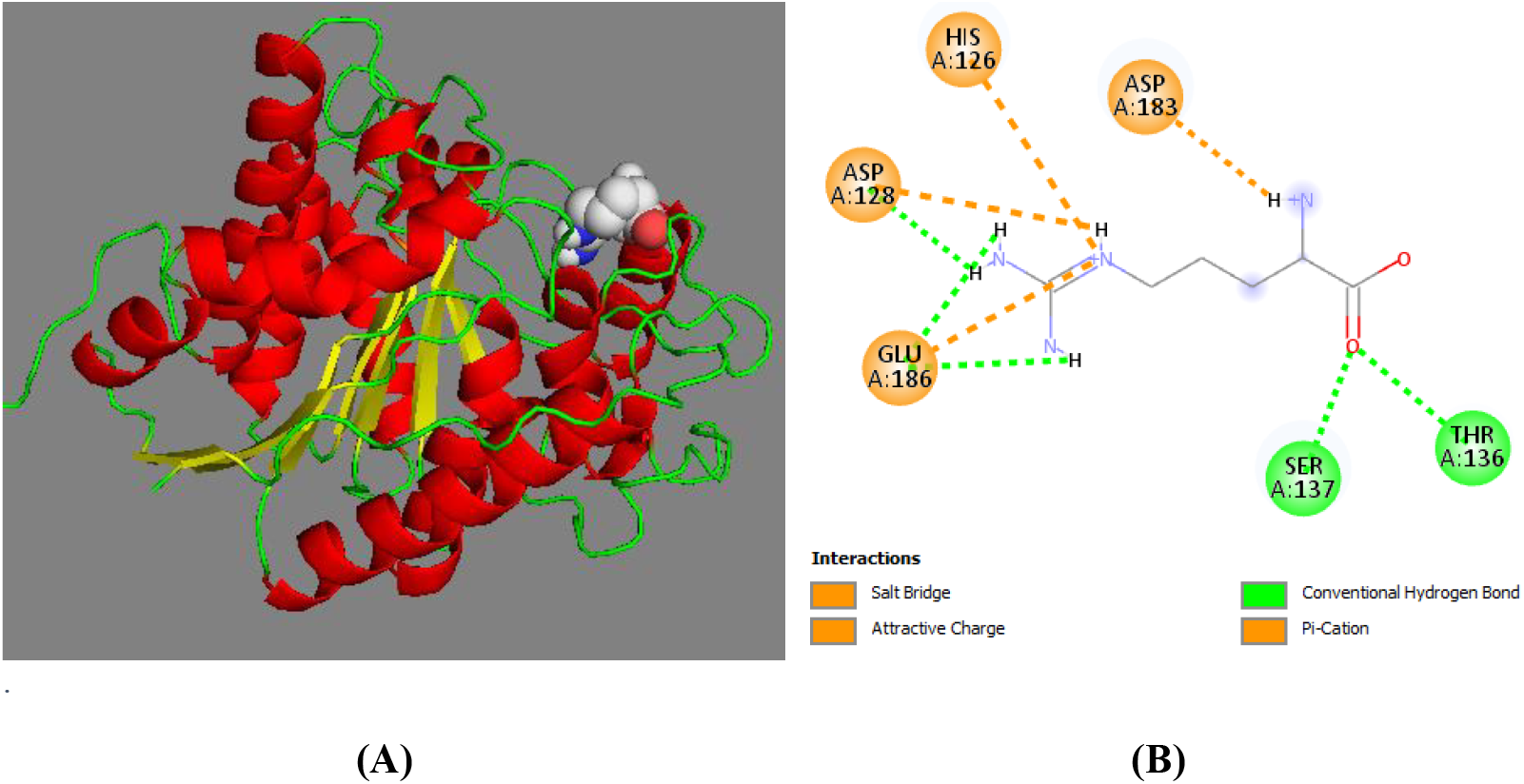
Elucidation of G119L–arginine interactions in 3D (A) and 2D (B) view.

### MD simulations

Data of MD simulations are shown in Fig. 5. The RMSD of the mutant form (i.e., the G119L variant) exhibited more harmonic overall protein motion compared to WT. In addition, the RMSD values of G119L were less than those of the WT, which indicates the greater stability of the mutant form. On the other hand, the WT of arginase manifested fewer fluctuations during a 50 ns simulation time than did the G119L mutant. The mutated Leu at 119 position had a very low RMSF value (0.0894 Å), which signifies the relaxed state of the mutant Leu at this position. The higher RMSF peaks can be traced to conformational changes that occurred to rearrange the interconnected AA in the tertiary structure in order to achieve a more relaxed state after mutagenesis. The two structures exhibited roughly similar compactness, as reflected by the convergent Rg. Likewise, the SASA of the WT had approximate values to the G119L variant (i.e., started from 144, 140 nm^2^ and ended at 131, 130 nm^2^). These findings mirror the stability, flexibility and relatively comparable conformations of the G119L mutant and the WT.

**Figure 5.**
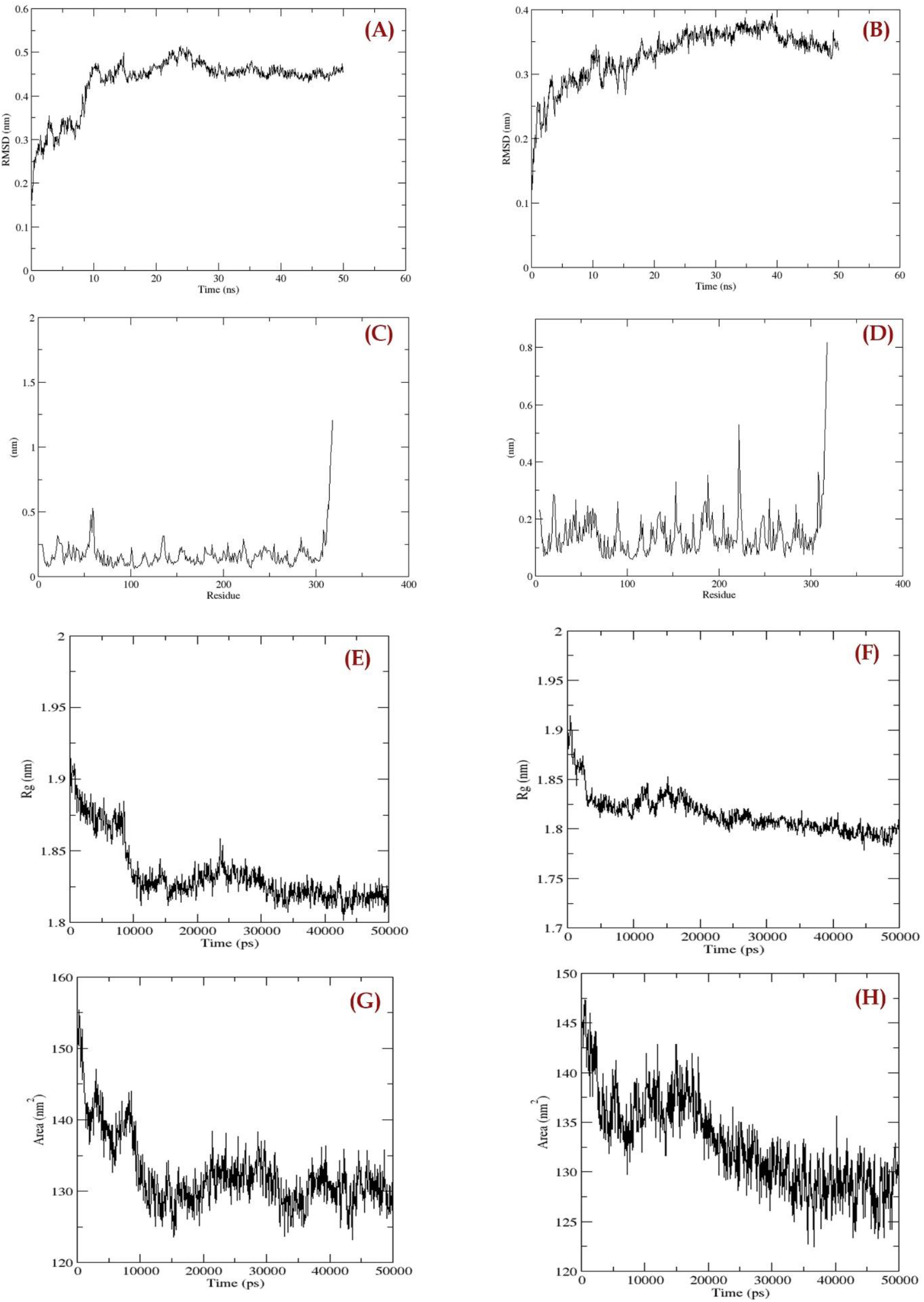
MD simulation of WT and G119L mutant for 50 ns. RMSD, RMSF, Rg and SASA for WT (A, C, E and G) and G119L mutant (B, D, F and H) are shown.

### In silico cloning

The CAI value of the back-translated sequence from the protein primary structure was 0.333, and the GC% was 66%. After the codon adaptation and optimisation conducted by the JCat tool, CAI and GC% values were substantially improved: 0.980 and 51%, respectively. The corresponding adaptiveness of the sequences are presented in Fig. 6.

**Figure 6.**
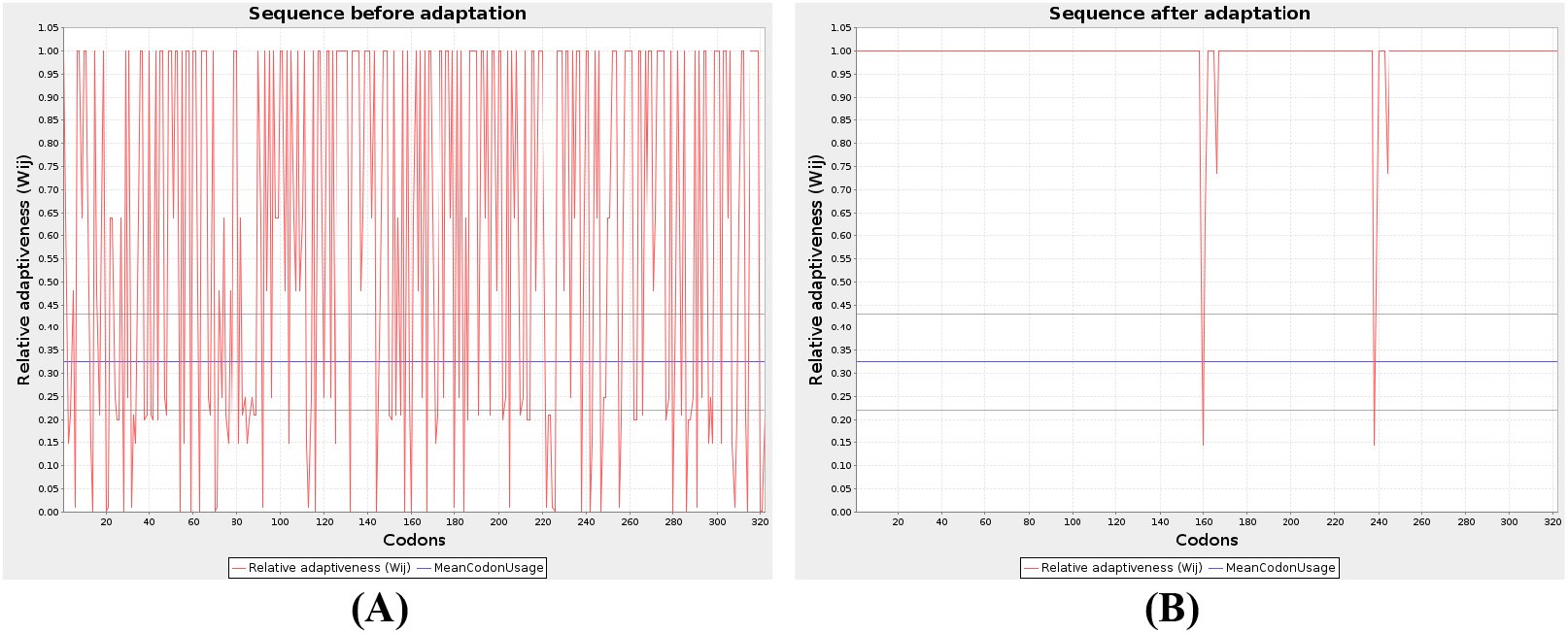
Relative adaptiveness of arginase back-translated sequence before (A) and after (B) Jcat treatment.

The codon-adopted sequence was 996 long nucleotides, and the whole vector was 6,353 bp. A 6X-His tag (i.e., six His residues) for affinity separation was embedded at the 3’ end of the inserted gene fragment. This sequence was inserted after introducing EcoRI and BamHI restriction sites at the beginning and end of the gene sequence. Restriction and insertion cloning were then performed between the corresponding restriction sites, as depicted in Fig. 7.

**Figure 7.**
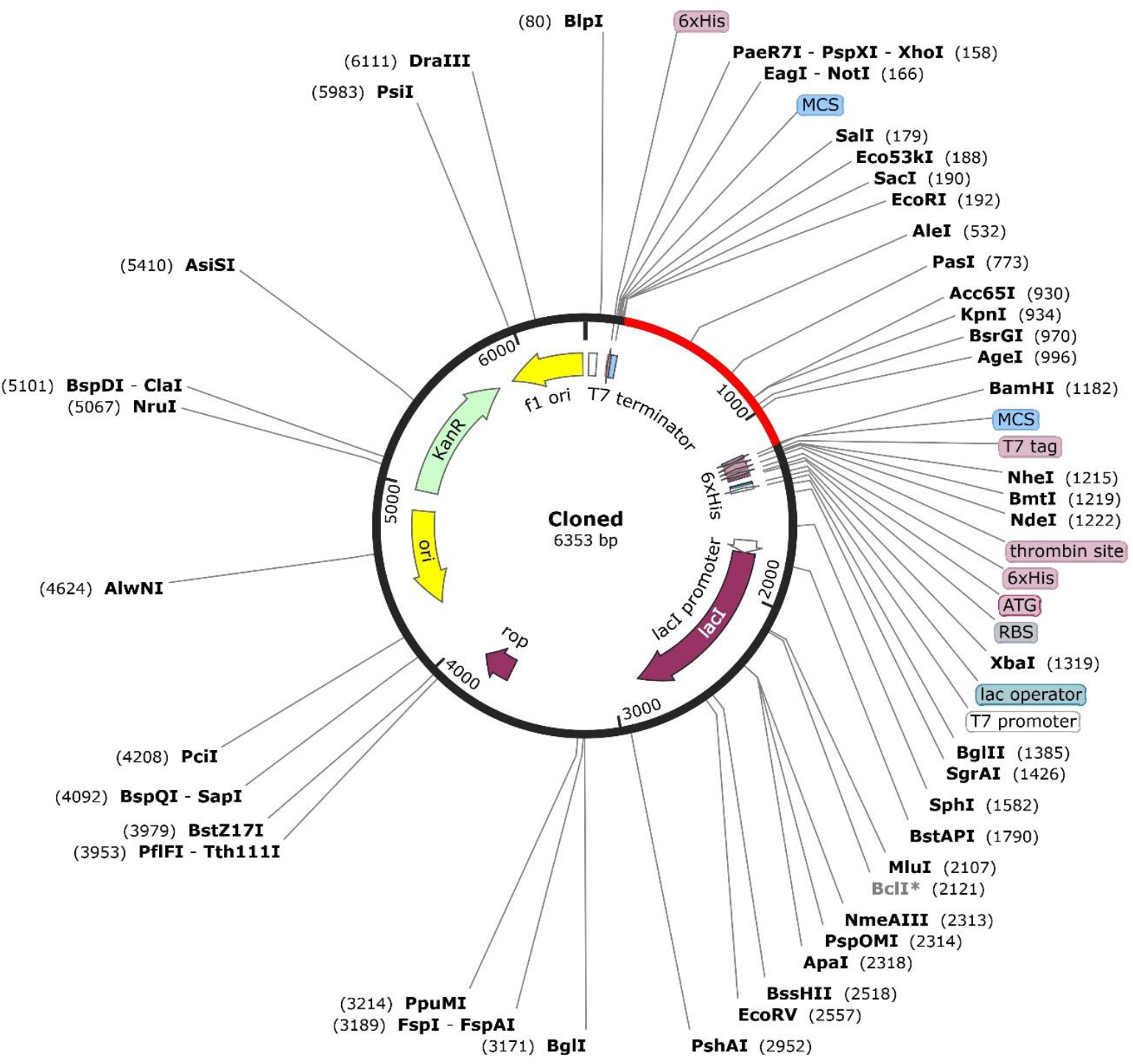
G119L-inserted pET-28(+)a plasmid; inserted sequence is highlighted in red.

### Protein-sol

The G119L mutant had an isoelectric point of 6.580, approaching neutral pH. Additionally, its solubility was estimated to be 0.521, which is greater than the solubility of average protein counterparts expressed in *E. coli* (Fig. 8); this facilitates the separation process from the other proteins expressed in *E. coli* and the His-tag added during cloning.

**Figure 8.**
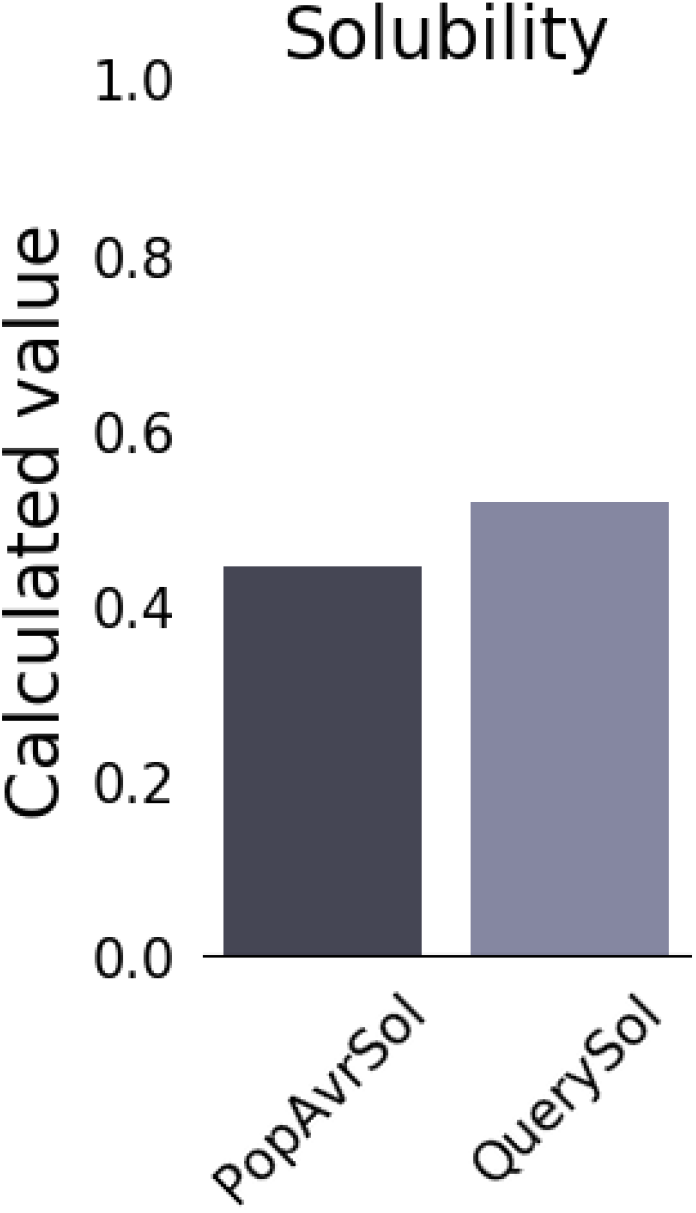
Predicted solubility of G119L mutant (QuerySol) in comparison to average proteins (PopAvrSol).

## Discussion

Arginase is a hydrolase enzyme capable of degrading arginine into urea and ornithine. This reaction is of great importance, rendering it a multifaceted enzyme that can be utilised for a variety of medical (i.e., to treat auxotrophic cancers [37] and arginase deficiency [38]) and industrial purposes (i.e., for manufacturing L-ornithine as a food additive). For arginase to satisfy these demands, it must have high activity and more stability to be recycled many times during the application process. Here is where the significance of the current study comes into play. The aim of this in silico work was to engineer the arginase enzyme to have a stronger binding affinity to the substrate, less inhibition and greater stability via various bioinformatics approaches.

The retrieved crystal structure of arginase from PDB demonstrated its high quality, as reflected by the low MolProbity score (1.21), high Ramachandran-favoured region occupancy and Z score of −8.79 in the ProsA tool; this enabled the commencement of silico mutagenesis. Among the examined AA residues that are located outside the binding pocket (in order to not interfere with the catalytic function of the enzyme), G119L was the best mutant with the highest binding affinity to the native substrate arginine (−5.9 kcal/mol), compared to the WT (−4.6 kcal/mol), with a 1.28-fold increase in theoretical activity.

Furthermore, G119L had weaker inhibition due to the reference inhibitor than WT (−5.4 vs. −5.6 kcal/mole). The strong binding is accounted for by multiple H-bonds, electrostatics and hydrophobic interactions formed to arginine after mutation. The introduction of Leu in place of Gly at 119 positions resulted in the formation of new H-bonds and hydrophobics near the active site, as Leu has a bulkier side chain than Gly. This explains the −0.12 kcal/mole increase in stability after mutagenesis given by the PremPS server output. Similarly, the RMSD, RMSF, Rg and SASA findings of the MD simulation for 50 ns reveals the stability, flexibility and comparable molecular motion of the G119L mutant to the WT enzyme. This confirmed the superior features of the mutant form, which instigated the provision of in-silico cloning as a preliminary step, paving the way for the applicability thereof in a wet-lab setting.

The protein sequence was back-translated into a nucleotide sequence, then optimised. The optimised sequence had a CAI value and GC% of 0.980 and 51%, compared to 0.333 and 65% prior to codon adaptation. The optimised sequence was then inserted into pET-28(+), a vector that is expressed in the *E. coli* K2 strain; pET-28a is one of the most popular expression vectors in the biotech market and is mentioned in more than 40,000 published articles. It has a copy number of 40, which indicates the high yield and ease of downstream purification of the recombinant protein [39]. An His-tag (6X-His) was inserted into the arginase sequence to ease its separation using Ni^+2^-affinity columns after the expression [40]. The solubility of the expressed protein was predicted to be 0.521, which reflects the potential high solubility since most *E. coli* expressed proteins had a value below 0.521 [36].

Reports regarding the engineering of arginase is too scarce which raises the significance of this work. Cheng et al. rationally screened variants of arginase in vitro for their potentiality to increase the polyethylene glycol (PEG) attachment on its surface to improve its enzyme stability *in vivo* [41]. Similarly, a novel single isoform of PEGylated human arginase was developed; this manifested potent cytotoxicity at sub-nM levels against cancer-cell lines of breast, prostate and pancreatic origin, due to arginine levels being diminished in a sustained manner [42]. These experimentally generated arginase variants transformed some of the AA residues that lie on the enzyme surface in order to enhance the stability with no goal of activity improvements. The findings of the present study therefore signify the novelty of generating such a variant by utilising the power of bioinformatics to conduct this exhaustive empirical work.

## Conclusion

The current work investigated the utilisation of various bioinformatics tools to generate engineered human arginase variants with stronger binding affinity to the substrate, less inhibition and greater stability. G119L was the best candidate to satisfy these criteria and exhibit an MD simulation pattern comparable to that of WT. In silico cloning of this variant was also provided to be expressed in *E. coli* K2, and the resulting protein was designed to have an His-tag for the facilitation of its downstream separation, since its solubility had higher-than-average the expression system proteins. The results of this theoretical work are recommended for testing in a wet-lab setting.

## Author contribution

Concepualiztion; methodology; data curation; drafting the manuscript: HA; investigation; supervision; editing and reviewing the manuscript: MA

